# Long noncoding RNA ANRIL supports proliferation of adult T-cell leukemia cells through cooperation with EZH2

**DOI:** 10.1101/329912

**Authors:** Zaowen Song, Wencai Wu, Mengyun Chen, Wenzhao Cheng, Juntao Yu, Jinyong Fang, Lingling Xu, Jun-ichirou Yasunaga, Masao Matsuoka, Tiejun Zhao

## Abstract

Adult T-cell leukemia (ATL) is a highly aggressive T-cell malignancy induced by human T-cell leukemia virus type 1 (HTLV-1) infection. Long noncoding RNA (lncRNA) plays a critical role in the development and progression of multiple human cancers. However, the function of lncRNA on HTLV-1-induced oncogenesis has not been elucidated. In the present study, we show that the expression of the lncRNA ANRIL was elevated in HTLV-1 infected cell lines and clinical ATL samples. E2F1 induced ANRIL transcription by enhancing its promoter activity. Knocking down of ANRIL in ATL cells repressed cellular proliferation and increased apoptosis *in vitro* and *in vivo*. As a mechanism for these actions, we found that ANRIL targeted EZH2, and activated the NF-κB pathway in ATL cells. This activation was independent of the histone methyltransferase (HMT) activity of EZH2, but required the formation of an ANRIL/EZH2/p65 ternary complex. Chromatin immunoprecipitation assay revealed that ANRIL/EZH2 enhanced p65 DNA binding capability. In addition, we observed that ANRIL/EZH2 complex repressed p21/CDKN1A transcription through H3K27 trimethylation of the p21/CDKN1A promoter. Taken together, our results implicate that lncRNA ANRIL, by cooperating with EZH2, supports the proliferation of HTLV-1 infected cells, which is thought to be critical for oncogenesis.

**IMPORTANCE:** Human T-cell leukemia virus type 1 (HTLV-1) is the pathogen that causes adult T-cell leukemia (ATL), which is a unique malignancy of CD4+ T cells. A role for long noncoding RNA (lncRNA) in HTLV-1-mediated cellular transformation has not been described. In this study, we demonstrated that lncRNA ANRIL was important for maintaining proliferation of ATL cells *in vitro* and *in vivo*. ANRIL was shown to activate NF-κB signaling through forming a ternary complex with EZH2 and p65. Further, epigenetic inactivation of p21/CDKN1A was involved in the oncogenic function of ANRIL. To the best of our knowledge, this is the first study to address the regulatory role of the lncRNA ANRIL in ATL and provides an important clue to prevent or treat HTLV-1 associated human diseases.

## INTRODUCTION

Human T-cell leukemia virus type 1 (HTLV-1) is a retrovirus that transforms human T lymphocytes(1). After a long latency period, about 2-5% of individuals infected with HTLV-1 develop adult T-cell leukemia (ATL)(1). HTLV-1 encodes several regulatory (*tax* and *rex*) and accessory (*p12*, *p13*, *p30*, and *HBZ*) genes in the pX region located between the env and 3’ long terminal repeat (LTR)(1). Among all of the HTLV transcripts, the *tax* gene is thought to play critical role in leukemogenesis because of its pleiotropic functions(2). Tax activates transcriptional pathways, including nuclear factor-κB (NF-κB), cAMP response element-binding protein (CREB), activator protein-1 (AP-1), and serum responsive factor (SRF)(2). The constitutive activation of the NF-κB pathway in HTLV-1 transformed T-cells argues for a critical role of this factor in mediating the development of ATL(3). However, Tax-mediated NF-κB activation may not fully explain ATL biology, because some leukemic cells that no longer express Tax continue to show constitutive NF-κB activation(4, 5). Recent reports provide new evidence that elevated expression of NF-κB-inducing kinase (NIK) has a pivotal role in the activation of the alternative NF-κB pathway in ATL independent of Tax expression(6). However, it remains unknown whether other mechanisms underlying the Tax-independent activation of the NF-κB pathway are involved in the development of ATL.

Regulatory noncoding RNAs (ncRNAs), such as microRNAs, small interfering RNAs, and long noncoding RNAs (lncRNAs), play important roles in the development of human diseases(7). LncRNAs, ranging from 200 to 100,000 nucleotides, are involved in a range of biologic processes, including modulation of cell growth, apoptosis, stem cell pluripotency, and immune response through the modulation of gene expression by epigenetic regulation, chromatin remodeling, transcription, and posttranscriptional processing(8, 9). Additionally, accumulating evidence has shown that lncRNAs play a critical role in tumorigenesis(10). However, the contribution by lncRNAs to the genesis of HTLV-1 induced ATL has not been investigated.

Recently, lncRNA ANRIL (antisense noncoding RNA in the INK4 locus), which is transcribed from the INK4b-ARF-INK4a gene cluster in the opposite direction, has been identified as a genetic susceptibility locus associated with human disease, in particular cancers(11-13). ANRIL was involved in repression of the p15/CDKN2B-p16/CDKN2A-p14/ARF gene cluster in *Cis* by directly binding to the polycomb repressor complex 2 (PRC2), which resulted in increased cell proliferation and suppression of apoptosis(14-16).

PRC2 has been demonstrated to be a functional target of some lncRNA, e.g., ANRIL, HOTAIR, Fendrr, H19, MALAT1, and COLDAIR(14, 15, 17-21). Moreover, PRC2/lncRNA complex-mediated dynamic control of H3K27 trimethylation (H3K27me3) is central to gene silencing in various cellular processes(8, 22). Enhancer of zeste homolog 2 (EZH2), one of the genes identified to be aberrantly overexpressed in ATL, is a component of PRC2. EZH2 contains a catalytic domain (SET domain) at the COOH-terminus that provides the methyltransferase activity, which plays a key role in the epigenetic maintenance of repressive chromatin marks(23, 24). In addition to its known role as transcriptional suppressor, several studies have also identified a PRC2-independent function of EZH2 in transcriptional activation rather than repression(25-28). In castration-resistant prostate cancer, EZH2 acts as a co-activator for critical transcription factors including the androgen receptor (AR)(29). This functional switch is dependent on phosphorylation of EZH2, and requires an intact methyltransferase domain. The activation role of EZH2 was also demonstrated in breast cancer cells, in which EZH2 activates NF-κB targets or NOTCH1(27, 28). However, the significance and potential role of polycomb group proteins and the associated lncRNA in ATL is still unknown.

In this study, we report that ANRIL interacted with EZH2 to support proliferation of ATL cells, indicating that dysregulation of the ANRIL is associated with the leukemogenesis of ATL.

## RESULTS

### LncRNA ANRIL is up-regulated in ATL

LncRNAs have been reported to be associated with the development of various cancers(7, 14, 19, 20). To identify lncRNAs that are involved in the development of ATL, we first examined the expression of selected onco-lncRNAs in HTLV-1–infected cell lines. Compared with HTLV-1 noninfected control cells, the level of ANRIL, H19, and SAF was enhanced in ATL cells, whereas three lncRNAs were found to be either slightly reduced (HOTAIR and TUSC7) or unchanged (MALAT1) in expression (Fig. S1). Since ANRIL was more highly expressed in ATL cells, our subsequent studies focused on ANRIL. The real-time PCR results verified the enhanced expression of ANRIL in HTLV-1-infected cell lines (Fig. 1A). Next, we examined the expression of ANRIL in primary ATL cells isolated from patients in comparison with PBMCs from healthy subjects under resting conditions. Primary ATL cells expressed ANRIL at levels much higher than healthy PBMCs (Fig. 1B).

**FIG 1.**
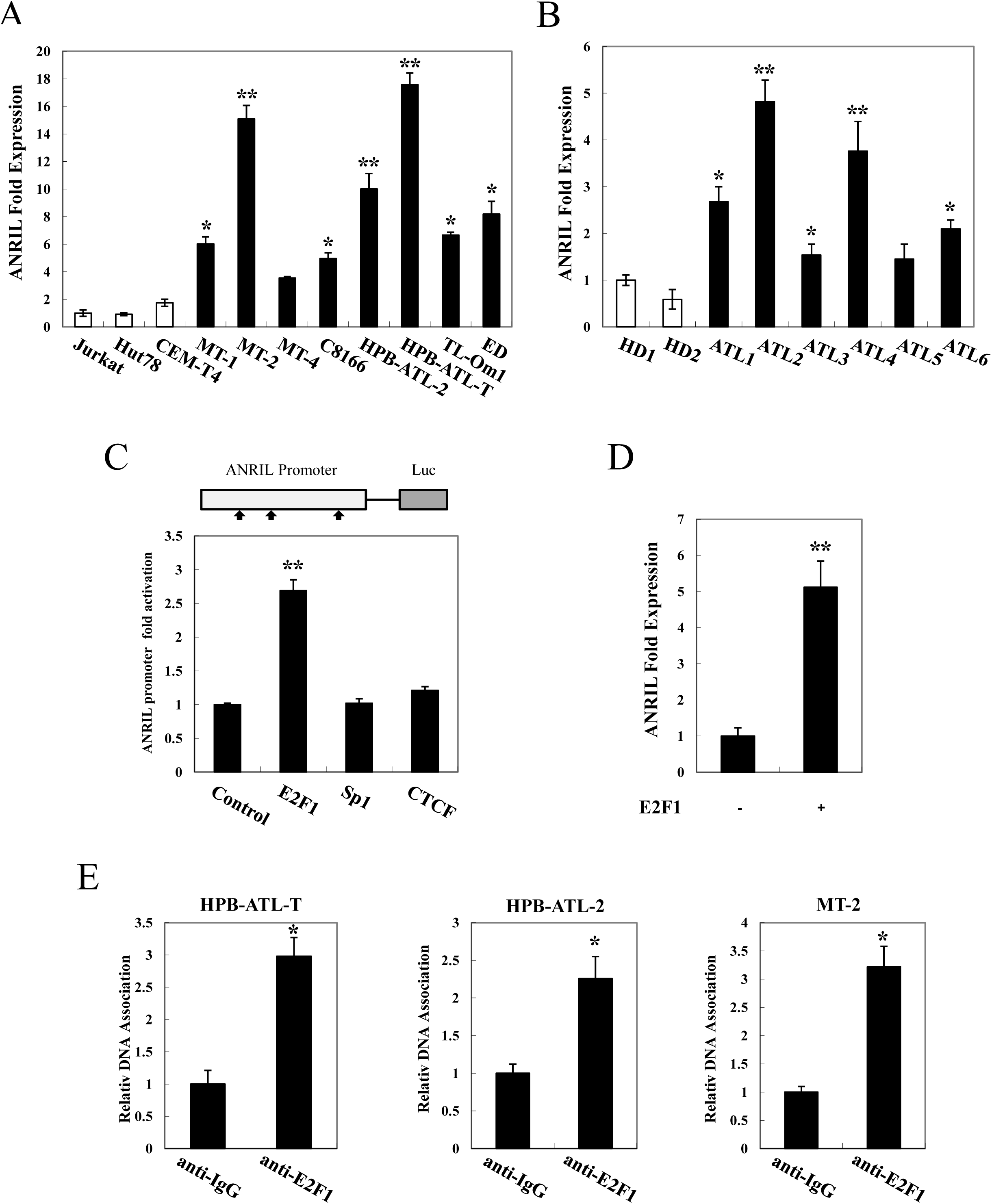
E2F1 induced high expression of lncRNA ANRIL in ATL. (A) lncRNA ANRIL is overexpressed in HTLV-1 associated cell lines. Total RNA was extracted from ATL and HTLV-I-immortalized cell lines (MT-1, MT-2, MT-4, C8166, HPB-ATL-2, HPB-ATL-T, TL-Om1, and ED) and T-cell lines not infected with HTLV-1 (Jurkat, Hut78, and CEM-T4)). The level of *ANRIL* mRNA was analyzed by quantitative PCR. (B) High expression of ANRIL in ATL. CD4 positive cells were isolated from PBMCs of healthy donors and ATL patients, and real-time PCR was performed to analyze the expression of ANRIL mRNA. HD1-2 indicates healthy donors, ATL1-6 indicates ATL patients. (C) E2F1 activated transcription of the ANRIL promoter. (Top panel) Schema of ANRIL promoter. The possible transcription factor binding sites are indicated. (bottom panel) 293FT cells were transfected with the E2F1 reporter plasmid with or without the expressing plasmids of E2F1, Sp1, and CTCF respectively. Luciferase activity was measured 48 hours after transfection. (D) E2F1 induced ANRIL expression. Total RNA was extracted from control or E2F1-expressing Jurkat cells. real-time PCR was performed to analyze the expression of *ANRIL* mRNA. (E) ChIP analysis of the association of endogenous E2F1 and ANRIL promoter in HPB-ATL-2, HPB-ATL-T, and MT-2 cells.

### E2F1 induces ARNIL expression in T cells

To investigate the mechanism of up-regulation of ANRIL in ATL cells, we amplified the putative promoter region of ANRIL (encompassing the −851 to +29 region), and cloned it into a pGL3-Basic reporter plasmid. Next, we searched for possible transcription factor binding sites in the promoter region using TESS (Transcription Element Search Software). E2F1, Sp1, and CTCF binding sites were found in this region. Reporter assay was performed to explore whether the putative transcription factors could affect ANRIL promoter activity. As shown in Fig. 1C, overexpression of E2F1 led to an increase of ANRIL (−851/+29)-luciferase expression, whereas Sp1 and CTCF had no effect. To investigate whether E2F1 indeed altered ANRIL expression in T cells, we over-expressed E2F1 in Jurkat cells. The results showed that enforced expression of E2F1 markedly upregulated the level of the ANRIL gene transcript (Fig. 1D). ChIP assays verified the binding of E2F1 to the promoter region of ANRIL in multiple ATL cell lines (Fig. 1E). These results collectively indicate that the enhanced expression of ANRIL in ATL can be attributed to the association of E2F1 with the ANRIL promoter.

### Knockdown of ANRIL impairs cellular proliferation and induces apoptosis *in vitro*

To clarify the potential role of upregulated ANRIL expression in ATL cells, we knocked down ANRIL by short hairpin RNA (shRNA) in HPB-ATL-T and ED cells. Two lentivirally expressed shRNAs directed against ANRIL resulted in a reduction of ANRIL expression compared with the vector control (Fig. 2A and B). MTT assays revealed that reduction of ANRIL expression significantly inhibited cell proliferation both in HPB-ATL-T and ED cell lines (Fig. 2C and D). In fact, ANRIL knockdown resulted in a 3-4 fold decrease in clonogenic survival of HPB-ATL-T cells (Fig. 2E). shANRIL#2 induced less-efficient knockdown of ANRIL than shANRIL#1, resulting in less inhibition of cell proliferation. In addition, ANRIL knockdown increased the total number of apoptotic cells in both the HPB-ATL-T and ED cell lines (Fig. 2F).

**FIG 2.**
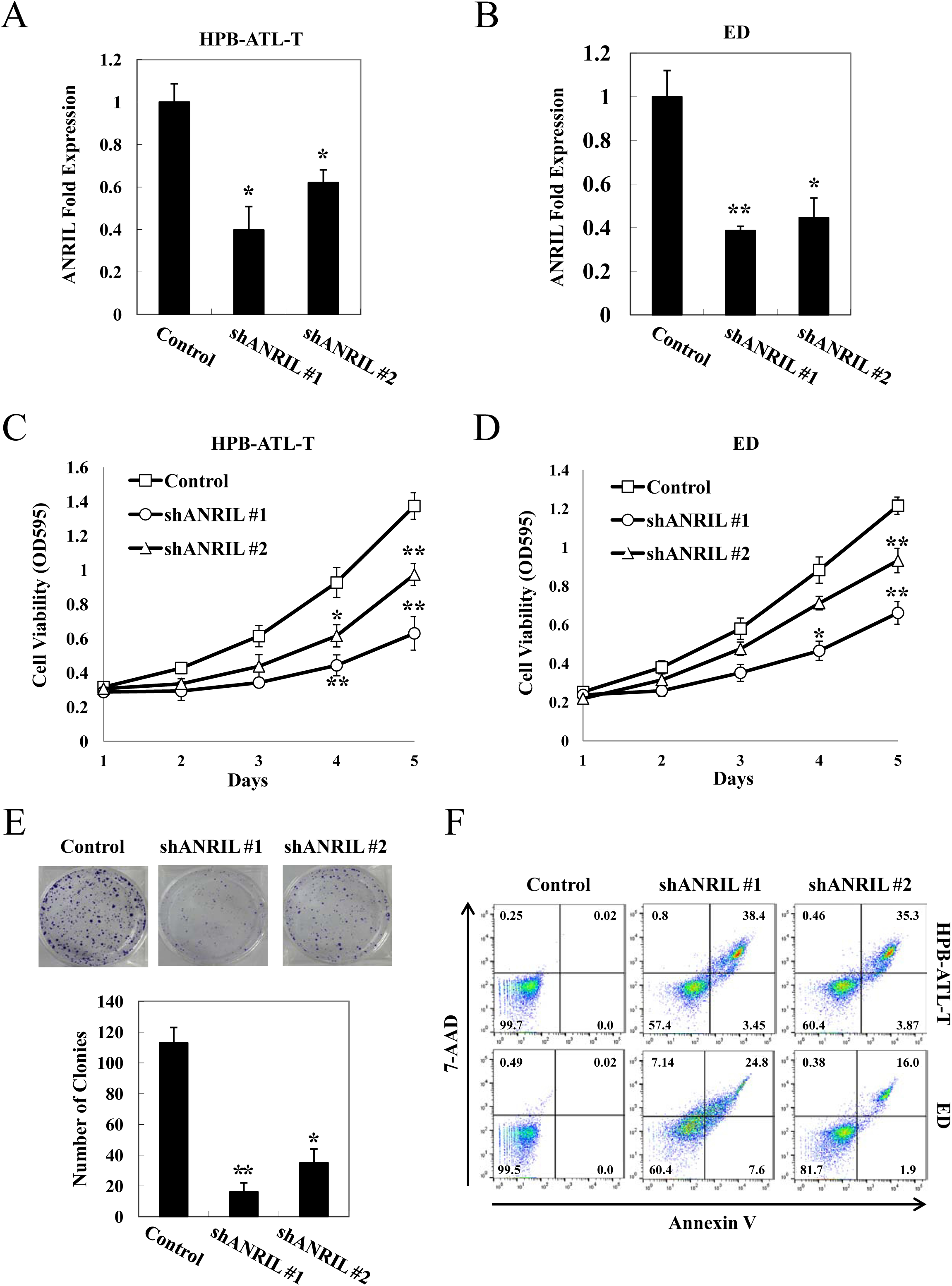
ANRIL supported proliferation of ATL cells. (A and B) Relative mRNA levels of ANRIL in HPB-ATL-T (A) and ED (B) cells transfected with recombinant lentivirus expressing pLKO-negative control (NC) or pLKO-shANRILs respectively. After puromycin selection, total RNA were extracted. The level of ANRIL mRNA was analyzed by real-time PCR. (C) Effect of ANRIL shRNAs on proliferation of HPB-ATL-T cells was determined by MTT assay. (D) MTT assay analysis of ANRIL shRNAs or mock-transfected ED cells. (E) Effect of ANRIL shRNAs on proliferation of HPB-ATL-T cells was determined by colony-formation assay. (F) Effect of ANRIL shRNAs on apoptosis of HPB-ATL-T and ED cells was analyzed by Annexin V and 7-AAD staining and flow cytometry method. The percentages of each fraction are shown. Statistical analysis was performed by Students *t* test with \**P*<0.05; \*\**P*<0.01.

Next, we evaluated if the effect of knockdown of ANRIL on ATL cell proliferation was due to cell cycle arrest. Flow cytometry results indicated that the proportion of cells in G1/S phase increased in ANRIL knockdown cells compared with the corresponding control. This suggested that reduction of ANRIL expression blocked cell cycle progression (Fig. S2).

Taken together, these observations demonstrate that ANRIL silencing suppressed proliferation of ATL cells, and this inhibitory effect was partially due to the inhibition of cell cycle progression and the promotion of apoptosis.

### ANRIL promotes proliferation of ATL cells *in vivo*

To further investigate the ability of ANRIL to stimulate ATL tumorigenesis *in vivo*, we injected an equivalent number of ED negative control and ED ANRIL knockdown cells into NCG mice and analyzed tumor growth. The tumors formed by ED-shANRIL#1 were smaller in both size and weight compared to ED-control tumors (Fig. 3A, B and C). Real-time PCR revealed that tumor nodules originating from ED-shANRIL#1 cells had decreased *ANRIL*, *MYC*, *VCAM1*, *CDK6*, and *CCND1* genes, and increased *CDKN1A* and *CTGF* expression compared to nodules originating from ED-control cells (Fig. 3D). H&E staining of the tumors showed that the degree of cell differentiation was significantly higher in ANRIL knockdown derived tumors compared to the control tumors (Fig. 3E).

**FIG 3.**
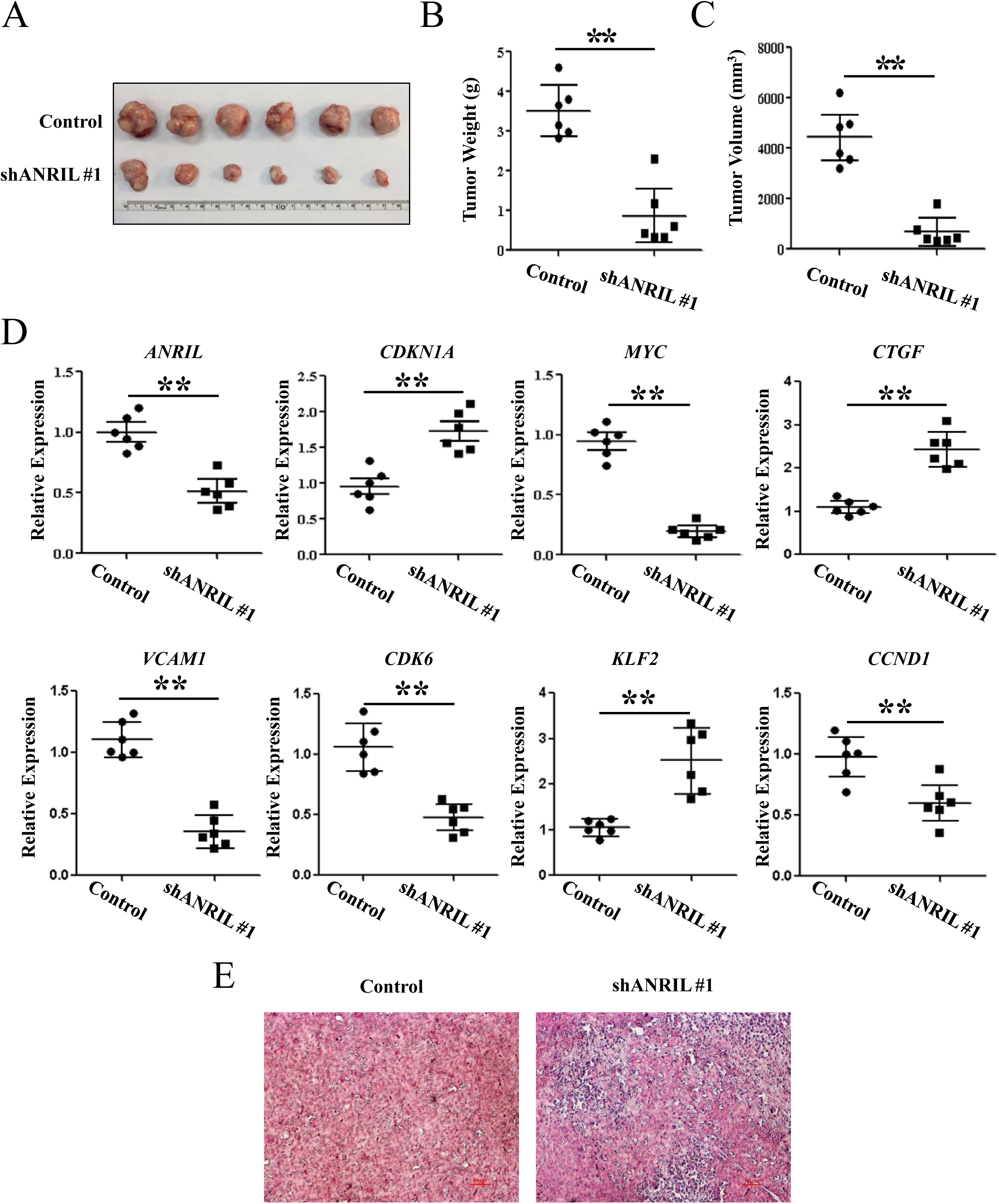
ANRIL promoted proliferation of ATL cells *in vivo*. Negative or ANRIL knockdown ED cells were injected into the NCG mice (n=6), respectively. 30 days after injection, all mice were sacrificed. (A) Photographs of dissected tumors from NCG mice. (B) Diagram of average weight of tumors. (C) Diagram of average volumes of tumors. (D) Expression levels of *ANRIL*, *CDKN1A*, *MYC*, *CTGF*, *VCAM1*, *CDK6*, *KLF2*, and *CCND1* were examined by real-time PCR in tumor tissues from NCG mice, respectively. (E) The tumor sections were under H&E staining. \**P*<0.05; \*\**P*<0.01.

Overall, these results confirm our *in vitro* transformation assays and suggest that ANRIL has tumor-forming potential.

### ANRIL and EZH2 synergistically activate the NF-κB pathway

Interestingly, as shown in Fig. 3D, the genes that were differentially expressed upon ANRIL depletion are many well-known NF-κB target genes, such as *MYC*, *VCAM1*, *CDK6*, and *CCND1*. Therefore, we evaluated whether ANRIL could influence the NF-κB pathway in ATL. As shown in Fig. 4A, silencing ANRIL expression in HPB-ATL-T and ED cell lines resulted in the suppression of NF-κB signaling. Reporter analyses revealed that the expression of ANRIL up-regulated Tax-mediated NF-κB activation. Treatment with SN50, an inhibitor of p65 activity, significantly suppressed the ability of Tax and ANRIL to enhance transcriptional activity through κB-responsive elements, indicating that ANRIL/Tax-mediated NF-κB activation is dependent on p65 (Fig. 4B).

**FIG 4.**
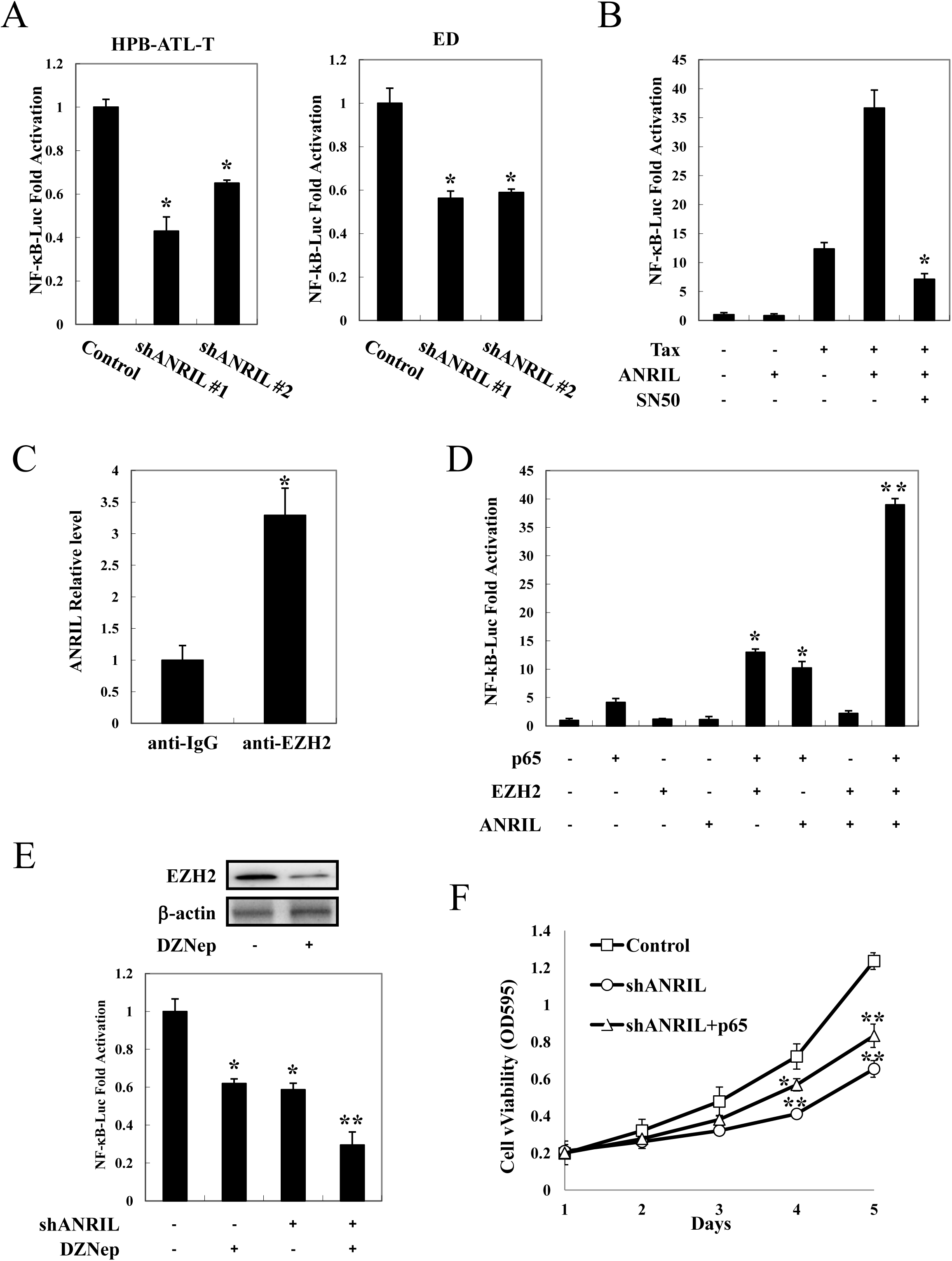
ANRIL/EZH2 activated NF-κB pathway. (A) Inhibition of NF-κB signaling by ANRIL knockdown. ANRIL knockdown HPB-ATL-T (left) and ED (right) cells were cotransfected with κB-Luc and phRL-TK plasmids. Luciferase levels were measured after 48 hours. (B) SN50 suppressed Tax and ANRIL induced NF-κB activation. Jurkat cells were cotransfected with κB-Luc, phRL-TK, pCG-Tax, and pCSII-CMV-ANRIL. Twenty-four hours after transfection, SN50 (20 μM) was added. After 24 hours, the cells were harvested and analyzed for luciferase activity. (C) ANRIL associated EZH2 *in vivo*. RIP analysis of the association of endogenous ANRIL RNA and EZH2 protein in HPB-ATL-T cells. (D) ANRIL and EZH2 synergistically reinforced p65-mediated NF-κB activation. Jurkat cells were cotransfected with κB-Luc, phRL-TK, pCSII-CMV-ANRIL, lenti-Myc-EZH2 and pCMV-p65. After 48 hours, the cells were harvested, and luciferase activity was determined. (E) DZNep treatment suppressed the NF-κB signaling in ANRIL knockdown cells. ANRIL knockdown cells were cotransfected with κB-Luc and phRL-TK plasmids. Twenty-four hours after transfection, DZNep (5 μM) was added. After 24 hours, the expression of EZH2 and β-actin were analyzed by immunoblotting (top panel), and the cells were harvested and analyzed for luciferase activity (bottom panel). (F) Over-expression of p65 rescued cell proliferation inhibition by ANRIL silencing. HPB-ATL-T cells were transfected with recombinant lentivirus expressing pLKO-shANRIL vector together with pCSII-CMV-p65. After puromycin selection, cell growth was detected by MTT assay. All the data shown are relative values of firefly luciferase normalized to Renilla luciferase and expressed as mean of a triplicate set of experiments (±SD). \**P*<0.05; \*\**P*<0.01.

It is well established that lncRNA may achieve regulatory goals through interacting with protein partners, and modulating the activity of these proteins within the cell. EZH2, which is a functional target of some lncRNAs, was constitutively expressed in all HTLV-1-infected cell lines and primary ATL cells (Fig. S3A and B). The association between ANRIL and EZH2 was confirmed to occur endogenously in an ATL cell line, HPB-ATL-T (Fig. 4C). Consistent with a previous report(27), EZH2 enhanced p65-induced NF-κB activation (Fig. S3C). Furthermore, in the presence of ANRIL, p65/EZH2-mediated NF-κB activation was dramatically reinforced, whereas it had a less obvious effect without EZH2 (Fig. 4D). To confirm the synergistic effect of ANRIL and EZH2 on NF-κB activation, we measured κB-Luciferase activity after suppressing EZH2 expression in ANRIL-silenced cells. Treatment with DZNep, an EZH2 inhibitor, further suppressed NF-κB signaling in ANRIL knockdown cells, indicating that ANRIL and EZH2 could cooperatively activate NF-κB pathway (Fig. 4E).

Next, we detected whether p65 was required for the ANRIL-enhanced cell proliferation of ATL cells. MTT assay showed that the decreased cell proliferation ability by shANRIL was partially rescued by p65 (Fig. 4F). These results together suggest that ANRIL supports proliferation of ATL cells through activating NF-κB pathway via EZH2.

### ANRIL associates with EZH2 and p65, and enhances NF-κB signaling independent of HMT activity

To clarify the mechanism by which ANRIL/EZH2 enhances NF-κB transcriptional response, FLAG-tagged p65 and myc-tagged EZH2 were cotransfected into 293FT cells. An interaction of EZH2 with p65 was demonstrated (Fig. 5A). Furthermore, ANRIL greatly enhanced the interaction between EZH2 and p65 (Fig. 5B). As ANRIL enhanced the activation of NF-κB signaling by EZH2 and p65, we explored whether ANRIL could form a complex with EZH2 and p65. 293FT cells were transfected with vectors expressing ANRIL, Myc-EZH2, and FLAG-p65, and a serial RNA immunoprecipitation assay was performed. As shown in Fig. 5C, we detected a specific ternary complex only when the three components were coexpressed. To investigate the binding of EZH2/ANRIL/p65 to DNA, we performed a ChIP assay in 293FT cells that were cotransfected with an κB-Luc reporter along with expression vectors for EZH2, ANRIL, and p65. The ChIP assay showed that EZH2 and ANRIL dramatically enhanced p65 DNA binding capability (Fig. 5D). Moreover, interaction of p65 to NF-κB binding site was diminished in ATL-T cells silencing both ANRIL and EZH2 (Fig. 5E). These results together suggest that ANRIL augments the interaction between EZH2 and p65, thereby potentiating NF-κB signaling.

**FIG 5.**
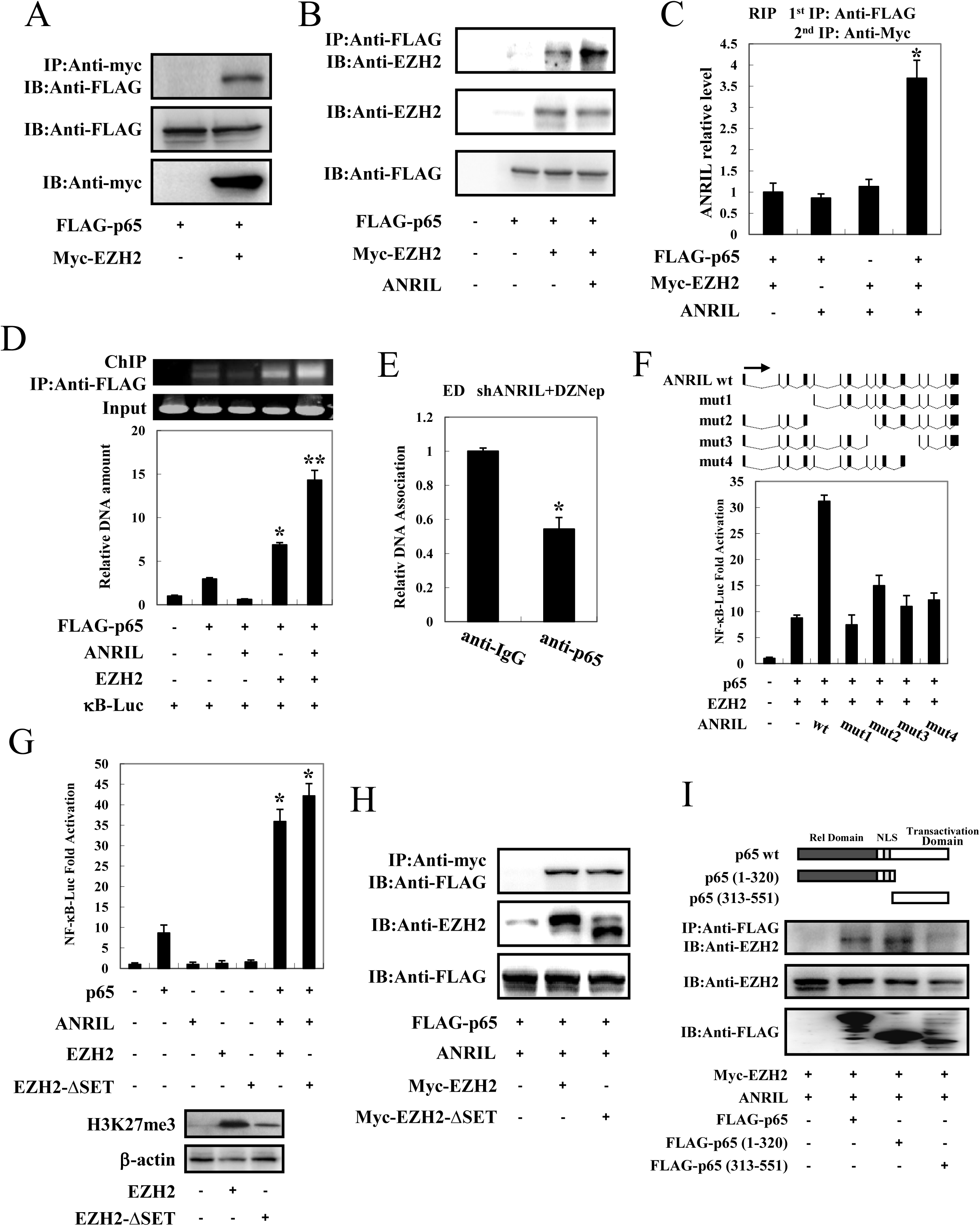
ANRIL formed complex with EZH2/p65 and enhanced NF-κB pathway independently of HMT activity. (A) EZH2 interacted with p65. 293FT cells were cotransfected with lenti-Myc-EZH2 together with FLAG-p65. After 48 hours, cell lysates were subjected to immunoprecipitation using anti–c-Myc followed by immunoblotting using anti-FLAG. (B) ANRIL enhanced the interaction between EZH2 and p65. 293FT cells were cotransfected with lenti-Myc-EZH2, pCSII-CMV-ANRIL, and pCMV-p65. Cell lysates were subjected to immunoprecipitation using anti-FLAG followed by immunoblotting with anti-EZH2. (C) ANRIL, EZH2 and p65 could form a ternary complex. pCSII-CMV-ANRIL, lenti-Myc-EZH2, and pCMV-p65 were cotransfected into 293FT cells. Ternary complexes were detected by sequential immunoprecipitation with anti-FLAG agarose affinity gel and anti-Myc antibody, and co-precipitated RNAs were detected by PCR. (D) ANRIL/EZH2 enhanced p65 DNA binding capability. After transfection with lenti-Myc-EZH2, pCSII-CMV-ANRIL, pCMV-p65, and KB-Luc for 48 hours, 293FT cells were chromatin immunoprecipitated by anti-FLAG antibody. The precipitated DNAs and 1% of the input cell lysates were amplified by the KB-Luc specific primers (top panel: RT-PCR, bottom panel: real-time PCR). (E) Silencing ANRIL and EZH2 inhibited p65 DNA binding capability. ANRIL knockdown cells were treated with DZNep. After 24 hours, the binding of p65 on DNA was analyzed by ChIP assay. (F) Analysis of ANRIL deletion mutants for their effect on p65/EZH2–induced NF-κB activation. Top panel: Schematic diagram of ANRIL and its mutants used in this study. Bottom panel: Jurkat cells were cotransfected with KB-Luc, phRL-TK, lenti-Myc-EZH2, pCMV-p65, and pCSII-CMV-ANRIL or its mutants. Luciferase activity was measured 48 hours after transfection. (G) ANRIL/EZH2 activated NF-κB pathway independent of EZH2 SET domain. Top panel: Jurkat cells were cotransfected with KB-Luc, phRL-TK, pCSII-CMV-ANRIL, pCMV-p65, together with lenti-Myc-EZH2 or lenti-Myc-EZH2ΔSET. At 48 hours after transfection, a dual luciferase reporter assay was performed. Bottom panel: The level of H3K27me3 was analyzed by immunoblotting. (H) ANRIL/EZH2 interacted with p65 independent of EZH2 SET domain. 293FT cells were transfected with expression vector of p65, pCSII-CMV-ANRIL, and lenti-Myc-EZH2 or lenti-Myc-EZH2ΔSET. After 48 hours, cell lysates were subjected to immunoprecipitation using anti–c-Myc followed by immunoblotting using anti-FLAG. (I) Mapping the region of the p65 protein necessary for the interaction with ANRIL/EZH2. Top panel: The schema of p65 deletion mutants has been shown. Bottom panel: 293FT cells were transfected with lenti-Myc-EZH2, pCSII-CMV-ANRIL, along with full-length or mutant FLAG-p65. At 48 hours after transfection, total cell lysates were subjected to IP using anti-FLAG followed by IB using anti-EZH2. \**P*<0.05; \*\**P*<0.01.

We next evaluated the ANRIL deletion mutants shown in Fig. 5F to determine which region of ANRIL is responsible for activating NF-κB signaling. Luciferase experiments revealed that the only full-length ANRIL could reinforced EZH2/p65-mediated NF-κB activation, whereas the truncated lncRNAs could not, suggesting that the tertiary structure of the full length of ANRIL, rather than one or more ANRIL domains was responsible for potentiating NF-κB signaling (Fig. 5F).

Accumulating evidence shows that the EZH2 SET domain, which has histone methyltransferase activity, is required to induce gene silencing by interacting with ANRIL(15, 21, 23, 24). We asked whether the HMT catalytic function of EZH2 is required for ANRIL/EZH2 complex to positively regulate NF-κB-dependent transcription in ATL cells. We first compared the EZH2 and EZH2 SET domain deletion mutant (EZH2-**Δ**SET) for the ability to activate the NF-κB pathway. As shown in Fig. 5G, lentiviral overexpression of EZH2-**Δ**SET resulted in a robust induction of NF-κB reporter activity, which was comparable to that of EZH2 wild type. In co-IP experiments in 293FT cells transfected with myc-tagged EZH2 or EZH2-**Δ**SET, we detected p65 in both EZH2 and EZH2-**Δ**SET immunoprecipitates (Fig. 5H). In addition, co-IP experiments revealed that ANRIL/EZH2 complex interacted with the Rel homology domain of p65 (Fig. 5I).

Taken all together, these data indicate that the complex of ANRIL/EZH2/p65 enhances NF-κB pathway in a histone methyltransferase activity independent manner.

### ANRIL/EZH2 suppresses p21/CDKN1A expression by histone methylation dependent mechanism

We next examined whether ANRIL support proliferation of ATL cells through modulating the expression of genes involved in apoptosis and proliferation. Cleaved caspase-3, caspase-7, and PARP were increased in HPB-ATL-T and ED cells with transfection of shANRIL (Fig. S4). Consistent with the results of the flow cytometry assay, in which knockdown of ANRIL induced cell cycle arrest, cyclin D1 and cyclin E1 expression was decreased in ANRIL knockdown HPB-ATL-T and ED cells (Fig. S4).

Several reports have indicated that ANRIL suppressed the expression of the p15/CDKN2B-p16/CDKN2A-p14/ARF gene cluster and contributed to cancer cell proliferation(14-16). However in the present study, there was no significant difference in the expression of p15/CDKN2B or p16/CDKN2A in ANRIL knockdown cells compared with control cells (Fig. 6A). Consistent with the observations *in vivo*, we did find that knockdown of the ANRIL gene led to increased p21/CDKN1A expression, indicating that p21/CDKN1A could be ANRIL novel targets in ATL (Fig. 6A).

**FIG 6.**
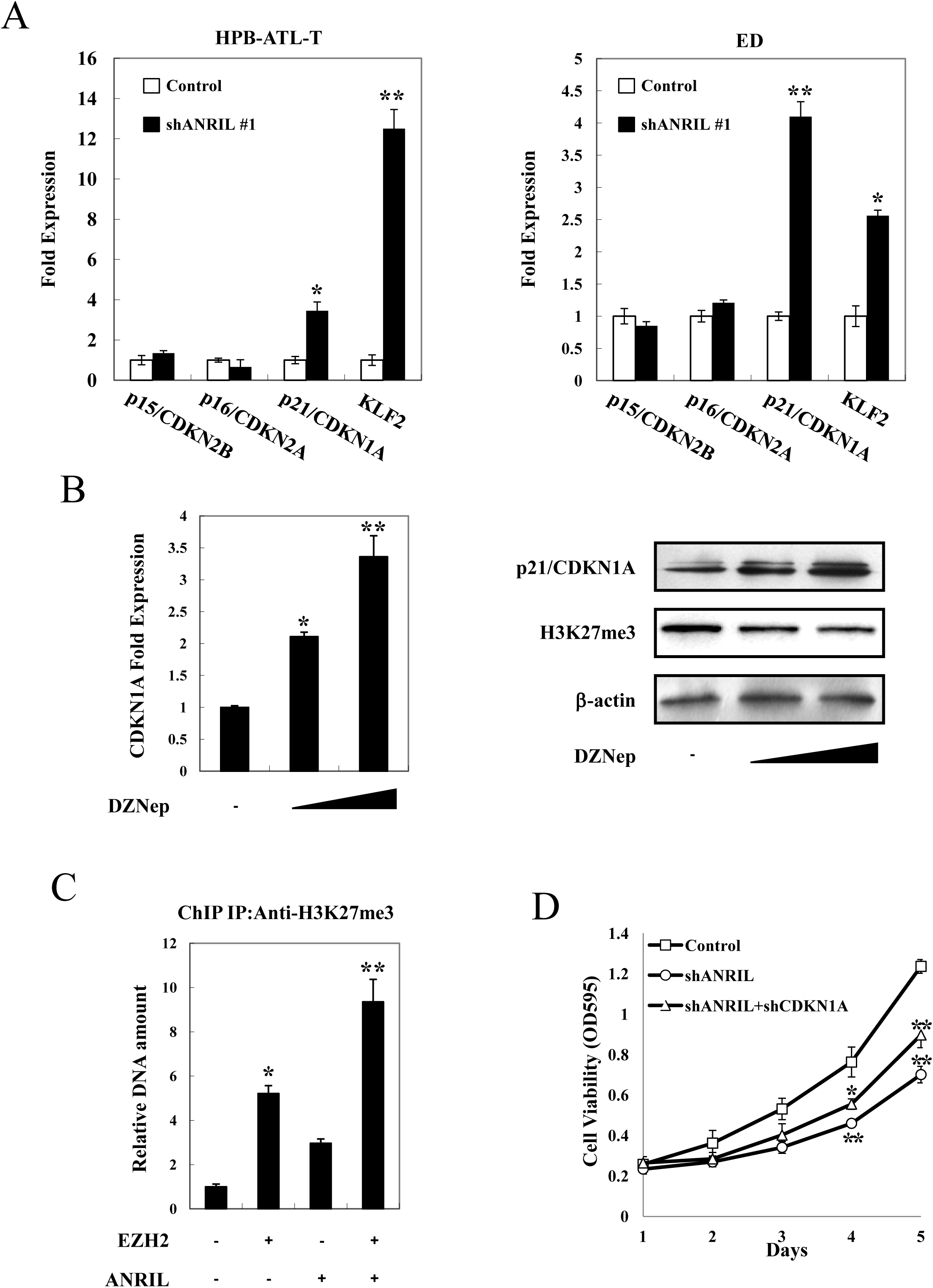
ANRIL/EZH2 suppresses p21/CDKN1A expression by histone methylation dependent mechanism. (A) Transcriptional changes of apoptosis and proliferation related genes in ANRIL knockdown HPB-ATL-T (left panel) and ED (right panel) cells. The levels of p15/CDKN2B, p16/CDKN2A, p21/CDKN1A, and KLF2 mRNA were analyzed by real-time PCR. (B) Treatment with DZNep induced the expression of p21/CDKN1A mRNA and protein. At 48 hours after DZNep treatment, the levels of p21/CDKN1A mRNA (left panel) and protein (right panel) were analyzed by real-time PCR and immunoblotting respectively. (C) ANRIL/EZH2 induced the accumulation of H3K27me3 on the p21/CDKN1A promoter. 293FT cells were transfected with lenti-Myc-EZH2 and pCSII-CMV-ANRIL. Forty-eight hours after transfection, 293FT cells were chromatin immunoprecipitated by H3K27me3 antibody. The precipitated DNAs and 1% of the input cell lysates were amplified by the specific primers for p21/CDKN1A promoter. (D) Down regulation of p21/CDKN1A rescued cell proliferation inhibition by ANRIL silencing. HPB-ATL-2 cells were transfected with recombinant lentivirus expressing pLKO-shANRIL vector together with pLKO-shCDKN1A. After puromycin selection, cell growth was detected by MTT assay. \**P*<0.05; \*\**P*<0.01.

It was previously reported that p21/CDKN1A was down regulated in ATL due to methylation of its promoter region(33). We next investigated if ANRIL could also affect the expression of p21/CDKN1A by histone methylation. After treatment with the EZH2 inhibitor DZNep, the mRNA and protein levels of p21/CDKN1A were increased in HPB-ATL-T cells (Fig. 6B). The results of a ChIP assay showed that overexpression of EZH2 resulted in the accumulation of H3K27 trimethylation on the promoter of p21/CDKN1A, which was further enhanced by ANRIL (Fig. 6C).

In order to evaluate the contribution of p21/CDKN1A to ANRIL-induced ATL cell proliferation, we performed a rescue experiment. As shown in Fig. 6D, down regulation of p21/CDKN1A in ANRIL silenced cells partially rescued shANRIL impaired proliferation of HPB-ATL-2 cells. These results suggest that ANRIL/EZH2 supports proliferation of ATL cells through inhibiting the expression of p21/CDKN1A via histone methylation dependent mechanism.

Collectively, these data support a model in which ANRIL/EZH2 complex not only contributes to the activation of NF-κB signaling, but also mediates transcriptional silencing of p21/CDKN1A in ATL. Moreover, we propose that the molecular mechanism through which ANRIL/EZH2 supports proliferation of ATL cells involves both histone methylation dependent and independent mechanisms (Fig. 7).

**FIG 7.**
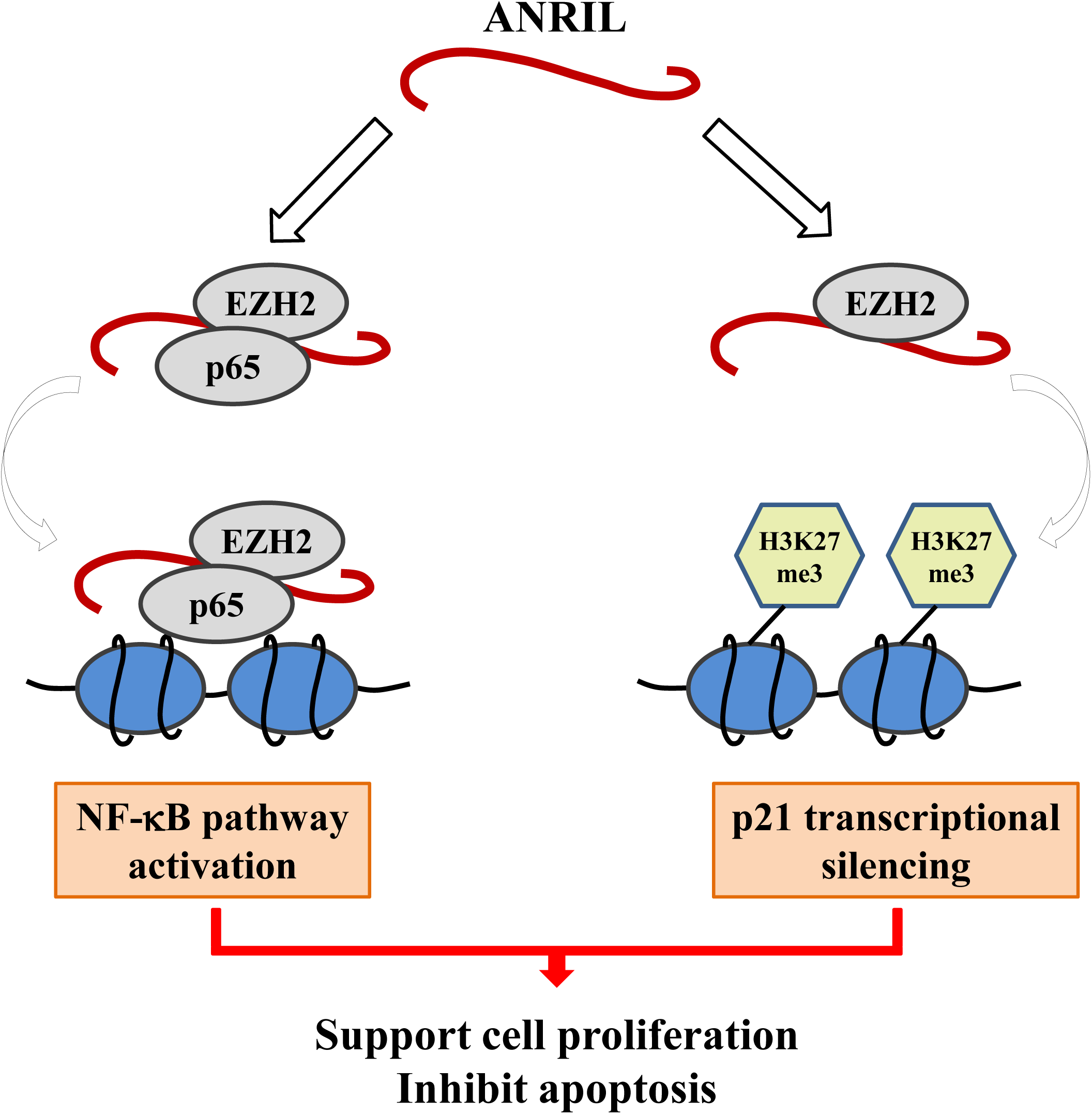
A schematic model that illustrates two functional arms of ANRIL in ATL. ANRIL targeted EZH2, and activated the NF-κB pathway via forming a ternary complex with p65. This activation was independent of EZH2’s histone methyltransferase activity. In addition, ANRIL/EZH2 complex inhibited the intrinsic apoptosis pathway through suppressing p21/CDKN1A in a methyltransferase activity-dependent manner.

## DISCUSSION

The NF-κB signaling pathway, constitutively activated in a variety of cancers, plays a major role in tumor biology as a regulator of key processes of cancer initiation and progression(34). In ATL, Tax-mediated NF-κB activation is reported to be critical for the proliferation of ATL cells and for the transforming activity of HTLV-1(35). However, ATL cells from most new cases, although infected with HTLV-1, are characterized by a loss of the expression of viral proteins, including Tax, because of host immune surveillance during the long latent period of the virus(1). The fact that NF-κB signaling is strongly and persistently activated in ATL cells implicates other viral and/or cellular factors. It has been reported that NIK may replace Tax in maintaining the constitutive activation of NF-κB, and NIK has been implicated in ATL pathogenesis(6). However, NIK does not seem to be essential for NF-κB activation in all ATL cells, as nuclear NF-κB was not exclusively expressed in NIK-overexpressing cell lines(6). EZH2, a histone methyltransferase subunit of a polycomb repressor complex, is highly expressed in a wide range of neoplasms, including cancers of the breast, lung, and pancreas, sarcomas, and lymphomas(36). The extensive evidence linking EZH2 activity to cancer has prompted interest in the underlying mechanism. We present evidence that EZH2 and its associated lncRNA ANRIL are highly expressed in ATL. EZH2 activated the NF-κB pathway in ATL cells through physical interaction with ANRIL and RelA (p65), and this positive regulation of NF-κB signaling did not require its methyltransferase activity. This constitutive activation by EZH2/ANRIL/RelA complex may explain how NF-κB is activated in the absence of Tax, and advances our understanding of how EZH2 may function as an oncogene. Consistent with our findings, it was recently reported that hepatitis B virus (HBV) infection induces EpCAM expression via RelA in complex with EZH2, and EZH2/RelA complex acts as an activator of NOTCH1 signaling to accelerate breast cancer growth, supporting that recruitment of RelA is essential for the oncogenic function of EZH2(28, 37).

Over-expression of an active form of EZH2 was demonstrated in ATL cells(38). Further study showed that the over-expressed EZH2 exhibited histone methyltransferase activity, as the ATL cells were strongly positive for H3K27me3(39). The EZH2-mediated global methylation gain at H3K27 caused silencing of tumor suppressors, transcription factors, and miRNAs, supporting leukemic cell characteristics. In addition to its known roles in histone modification and transcriptional regulation, EZH2 also promotes tumorigenicity of various cancers independent of its H3K27 methyltransferase activity, and of the polycomb repressive complex 2, through transcriptional activation marks(25-28). Our study defines a unique role and mechanism for EZH2 in ATL, whereby EZH2 utilizes polycomb-dependent and -independent mechanisms to protect ATL cells from apoptosis and promote their proliferation.

Recently, Yamagishi et al. reported that overexpression of EZH2 led to polycomb-mediated silencing of miR-31 and constitutive activation of NF-κB signaling(40). Our results complement this finding and highlight a novel mechanism for NF-κB activation in ATL, where an aberrant overexpression of EZH2 activates NF-κB through both HMT-dependent and -independent pathways. In addition, because EZH2 was undetectable in cells from healthy adults and HTLV-1 carriers, it is likely that deregulation of NF-κB caused by EZH2 is involved in the early steps of ATL oncogenesis. Association between EZH2 and RelA are essential for NF-κB activation, whereas the epigenetic modification function of EZH2 requires the recruitment of PRC2 subunits, such as SUZ12 and EED. Thus, we speculate that the bimodal function of the EZH2-ANRIL complex depends on the EZH2 binding partner.

Dysregulation of the lncRNA and EZH2 complex has been implicated in viral transformation. For example, lncRNA-HEIH was found to be involved in HBV-related hepato-carcinogenesis through association with the enhancer of EZH2, resulting in the promotion of hepatoma cell proliferation(41). HBx-upregulated lncRNA-UCA1 promotes cell growth and tumorigenesis by recruiting EZH2 and repressing p27Kip1/CDK2 signaling(42). In Kaposi’s sarcoma-associated herpesvirus, lncRNA-PAN RNA binds to protein components of polycomb repression complex 2, EZH2 and SUZ12, and globally influences cellular gene expression, leading to a complete abrogation of the initiation of the viral lytic-phase transcription program(43). Similarly, HIV-expressed lncRNA recruit and guide a chromatin-remodeling complex (consisting of proteins such as EZH2 and DNMT3a) to the viral promoter, resulting in reduced levels of viral gene expression(44). These findings show that dysregulation of lncRNA-EZH2 complex are common among different viruses, suggesting that these activities are critical for viral oncogenesis. To our knowledge, all reported instances of virus-induced dysregulation of the lncRNA-EZH2 complex resulted in modulation of global gene expression due to H3K27 methylation. Here we demonstrate that EZH2-ANRIL complex employs novel HMT-dependent and -independent mechanisms to support proliferation of ATL cells.

Global changes in the epigenetic landscape are a hallmark of cancer(45). Since genetic changes in specific genes in ATL cells have not been identified except for p53 and p16, and there is no consistent chromosomal change, it is possible that epigenetic changes such as DNA methylation, histone modification, and ncRNAs, which dysregulate transcription of oncogenes, play an important role in oncogenesis(46). In ATL cells, DNA methylation of the promoter regions of tumor suppressor genes (p21 and p16, etc.) and of the 5’LTR of HTLV-1 is commonly observed and accumulates with disease progression(1, 33, 47). A recent study showed that PRC2-mediated trimethylation at H3K27 (H3K27me3) was significantly and frequently reprogrammed at half of genes in ATL cells(39). In this study, we identify a novel mechanism of down regulation of p21/CDKN1A due to EZH2/ANRIL triggered histone H3K27 promoter trimethylation. It has recently become apparent that DNA methylation and histone modification pathways can be dependent on one another(48). Moreover, EZH2 controls CpG methylation through direct physical contact with DNA methyltransferases(49). It is thus likely that, apart from its regulatory function on p16/CDKN1A expression, EZH2 may dysregulate transcription of other tumor suppresser genes in ATL cells by two dependent epigenetic means to acquire malignant phenotypes. However, it remains unknown how EZH2 selectively regulates its target genes by DNA methylation or histone modification. We speculate that the regulatory effect of EZH2 on its target gene expression is dependent on its affinity to DNA, and may also depend on its associating transcription factors. Further studies are needed to fully elucidate the role of EZH2/ANRIL in HTLV-1 infection and ATL.

In summary, we showed that ANRIL supported the proliferation of ATL cells by interacting with EZH2. HTLV-1 may take advantage of these mechanisms to allow the infected cells to proliferate *in vivo*. These findings may contribute to the development of novel therapeutic strategies.

## MATERIALS AND METHODS

### Cell culture and clinical samples

HTLV-1-infected T-cell lines (MT-2, MT-4, C8166, MT-1, HPB-ATL-2, HPB-ATL-T, ED, and TL-Om1), and HTLV-1-negative T-cell lines (Jurkat, Hut78, and CEM-T4) were grown in RPMI 1640 supplemented with 10% fetal bovine serum (FBS) and antibiotics. 293FT cells were cultured in Dulbecco modified Eagle medium (DMEM) supplemented with 10% FBS and 500 μg/mL G418. Peripheral blood mononuclear cells (PBMCs) were isolated from ATL patients (n=6) and healthy volunteers (n=2). Details of clinical samples are shown in Table S1. Clinical samples were obtained and used according to the principles expressed in the Declaration of Helsinki and approved by the Institutional Review Board of Kyoto University. All patients provided written informed consent for the collection and use of samples.

### Plasmids and reagents

The sequence of ANRIL was amplified by reverse transcriptase-polymerase chain reaction (RT-PCR) and subcloned into pCSII-CMV-GFP. The ANRIL promoter sequence was cloned into pGL3-Basic vector (pGL3-ANRIL-851/+29). Expression vectors for E2F1, Sp1, Tax, p65, p65 deletion mutants, CTCF, κB-Luc, and phRL-TK were prepared as previously described(30, 31). Lenti-Myc-EZH2 plasmid was a gift from Dr Xiangping Li of Southern Medical University (China)(32). Expression vectors for Lenti-Myc-EZH2-**Δ**SET deletion mutants and pCSII-CMV-ANRIL deletion mutants were generated by PCR. 3-Deazaneplanocin A (DZNep) was purchased from Selleck (Houston, TX, USA). SN50 was purchased from Millipore (Billerica, MA, USA).

### Quantitative and semiquantitative PCR

Total RNA was isolated using Trizol Reagent (Invitrogen, Carlsbad, CA, USA) according to the manufacturer’s instructions. Complementary DNA was synthesized using the SuperScript III reverse transcriptase (Life Technologies, Grand Island, NY, USA). Quantitative PCR (qPCR) was carried out using Power SYBR Green PCR Master Mix and StepOnePlus Real Time PCR System (Thermo Fisher Scientific, Waltham, MA, USA). Semiquantitative RT-PCR was performed as previously described(30). The specific primers used can be found in Table S2.

### Luciferase assay

Jurkat cells were plated on 12-well plates at 1.2×10^5^ cells per well. After 24 hours, cells were transfected with the indicated plasmids using Lipofectamin LTX Reagent (Invitrogen). At 48 hours after transfection, a luciferase reporter assay was performed using the Dual-Luciferase Reporter Assay System (Promega, Madison, WI). Each experiment was performed in triplicate, and the data represent the mean and SD of 3 independent experiments, each normalized to Renilla activity.

### Chromatin immunoprecipitation assay

293FT cells were transfected with the indicated plasmids. At 48 hours after transfection, a ChIP assay was done according to the protocol recommended by Upstate Biotechnology. Precipitated DNA was amplified by PCR using primers specific for the ANRIL-Luc, κB-Luc, or p21/CDKN1A promoter vectors respectively. For the endogenous *ANRIL* promoter, chromatin samples prepared from ATL-T cells were subjected to ChIP analysis using anti-E2F1 (Abcam, Cambridge, MA) and normal mouse immunoglobulin G (Santa Cruz Biotechnology). Sequences for the primer set can be found in Table S2.

### Knockdown

Knockdown of ANRIL and p21/CDKN1A in ATL cell lines was performed with a lentiviral vector pLKO.1-based shRNA system (Open Biosystems, Lafayette, CO, USA) (31). Stable transfectants were selected in puromycin. The shRNA sequence specifically targeting ANRIL and CDKN1A was shANRIL #1: 5’-GGAATGAGGAGCACAGTGA-3’, shANRIL #2: 5’-AAGCGAGGTCATCTCATTGCTCTAT-3’, shCDKN1A: 5’-GCGATGGAACTTCGACTTT-3’.

### Cell proliferation assay

Cells were seeded in 96-well plates and at indicated time points, 10 μl of 3-(4,5-dimethythiazol-2-yl)-2,5-diphenyl tetrazolium bromide solution was added. Cells were incubated at 37 °C for 2 h and then lysed by 100 μl of lysis buffer (4% Triton X-100 and 0.14% HCl in 2-propanol). Absorbance at 595nm was measured on a Microplate Reader (Bio-Rad, Hercules, CA).

### Colony formation assay

Stable knockdown cells were plated on 6-well plates at 1000 cells per well, and kept in complete medium for 2 weeks. Colonies were fixed with methanol and stained with methylene blue. Colony formation was determined by counting the number of stained colonies. Triplicate wells were measured in each group.

### Measurement of apoptotic cell death

Apoptotic cells were routinely identified by Annexin V (Biolegend, San Diego, CA), and 7-amino-actinomycin D (7-AAD; BD Biosciences, San Jose, CA)-staining according to the manufacturer’s instructions and analyzed by flow cytometry (BD FACSVerse; BD Biosciences). Data files were analyzed by using FlowJo software (TreeStar, San Carlos, CA).

### *In vivo* tumorigenicity assay

Cell suspensions (4 × 10^6^ cells) was injected subcutaneously into the right flank of (NOD-Prkdc^em26Cd52^Il2rg^em26Cd22^/Nju, female NCG mice NOD/ShiLtJNju based immune deficient mice) (Model Animal Research Center of Nanjing University, China). Thirty days after injection, all mice were killed. Tumors weight and the expression of ANRIL RNA and its target genes were measured. All animals used in this study were maintained and handled according to protocols approved by Zhejiang Normal University and Huaqiao University.

### Histological analyses

Tissue samples were fixed in 10% formalin in phosphate buffer and then embedded in paraffin. Haematoxylin and eosin (H&E) staining was performed according to standard procedures. Images were captured using confocal laser scanning microscope (Leica TCS SP5 AOBS).

### Immunoprecipitation (IP) and immunoblotting

293FT cells were transfected with the indicated plasmids by Lipofectamine 3000 Reagent (Invitrogen). Tagged proteins were isolated from transfected 293FT cells by immunoprecipitation with anti–c-Myc (clone 9E10, Sigma-Aldrich, St Louis, MO) or anti-FLAG M2 (Sigma-Aldrich) antibodies, and analyzed by immunoblotting as described previously. Other antibodies used were as follows: anti–mouse Immunoglobulin G (IgG) and anti–rabbit IgG (GE Healthcare Life Sciences); anti-EZH2, anti-p65, anti-Caspase 3, anti-Caspase 7, anti-Caspase 8, anti-PARP, anti-p21, anti-Cyclin D1, and anti-Cylin E1 (Cell Signaling Technology, Danvers, MA, USA); anti-Trimethyl-Histone H3 (Lys27) (Millipore); anti-HA (Sigma-Aldrich); anti-β-actin (Bioworld, St. Louis Park, MN, USA).

### RNA immunoprecipitation

RNA immunoprecipitation was performed using a MagnaRIP RNA-binding protein immunoprecipitation kit (Millipore) according to the manufacturer’s instructions. For sequential RNA immunoprecipitation, transfected 293FT cells were lysed in radioimmunoprecipitation assay buffer and immunoprecipitated with anti-FLAG M2 agarose affinity gel. The precipitates were eluted with FLAG peptide and the eluate diluted with radioimmunoprecipitation assay buffer, immunoprecipitated with anti-Myc antibody, and co-precipitated RNAs were detected by PCR. The EZH2 antibody used for RNA immunoprecipitation was purchased from Cell Signaling Technology. The primers used to detect ANRIL expression can be found in Table S2.

### Statistical analyses

Statistical analyses were performed using the unpaired Student *t* test. Significance was set at alpha = 0.05.

## ACKNOWLEDGMENTS

We thank Caryl Antalis for proof-reading of this manuscript.

## Author contributions

Z.S., W.W., M.C., and T.Z. designed the research; Z.S., W.W., M.C.,W.C., J.Y., J.F. and T.Z. performed the research; J-I.Y. and M.M. provided samples and vital reagents; Z.S., W.W., M.C.,W.C., J.Y., L.X. and T.Z. analyzed the data; Z.S., W.W., M.C., and T.Z. wrote the paper.

